# Electrospun Fiber Surface Roughness Modulates Human Monocyte-Derived Macrophage Phenotype

**DOI:** 10.1101/2024.08.30.610568

**Authors:** Aidan Alemifar, KaLia Burnette, Bryan Jandres, Samuel Hurt, Hubert M. Tse, Jennifer L. Robinson

**Affiliations:** Department of Orthopaedic Surgery and Sports Medicine, University of Kansas Medical Center; Department of Mechanical Engineering, University of Kansas Medical Center; Department of Biochemistry, University of Washington, University of Kansas Medical Center; Bioengineering Graduate Program, University of Kansas Medical Center; Department of Chemical and Petroleum Engineering, University of Kansas, University of Kansas Medical Center; Department of Microbiology, Molecular Genetics and Immunology, University of Kansas Medical Center

## Abstract

Injuries to fibrous connective tissues have very little capacity for self-renewal and exhibit poor healing after injury. Phenotypic shifts in macrophages play a vital role in mediating the healing response, creating an opportunity to design immunomodulatory biomaterials which control macrophage polarization and promote regeneration. In this study, electrospun poly(-caprolactone) fibers with increasing surface roughness (SR) were produced by increasing relative humidity and inducing vapor-induced phase separation during the electrospinning process. The impact of surface roughness on macrophage phenotype was assessed using human monocyte-derived macrophages *in vitro* and *in vivo* using B6.Cg-Tg(Csf1r-EGFP)1Hume/J (MacGreen) mice. *In vitro* experiments showed that macrophages cultured on mesh with increasing SR exhibited decreased release of both pro- and anti-inflammatory cytokines potentially driven by increased protein adsorption and biophysical impacts on the cells. Further, increasing SR led to an increase in the expression of the pro-regenerative cell surface marker CD206 relative to the pro-inflammatory marker CD80. Mesh with increasing SR were implanted subcutaneously in MacGreen mice, again showing an increase in the ratio of cells expressing CD206 to those expressing CD80 visualized by immunofluorescence. SR on implanted biomaterials is sufficient to drive macrophage polarization, demonstrating a simple feature to include in biomaterial design to control innate immunity.

**GRAPHICAL ABSTRACT:** 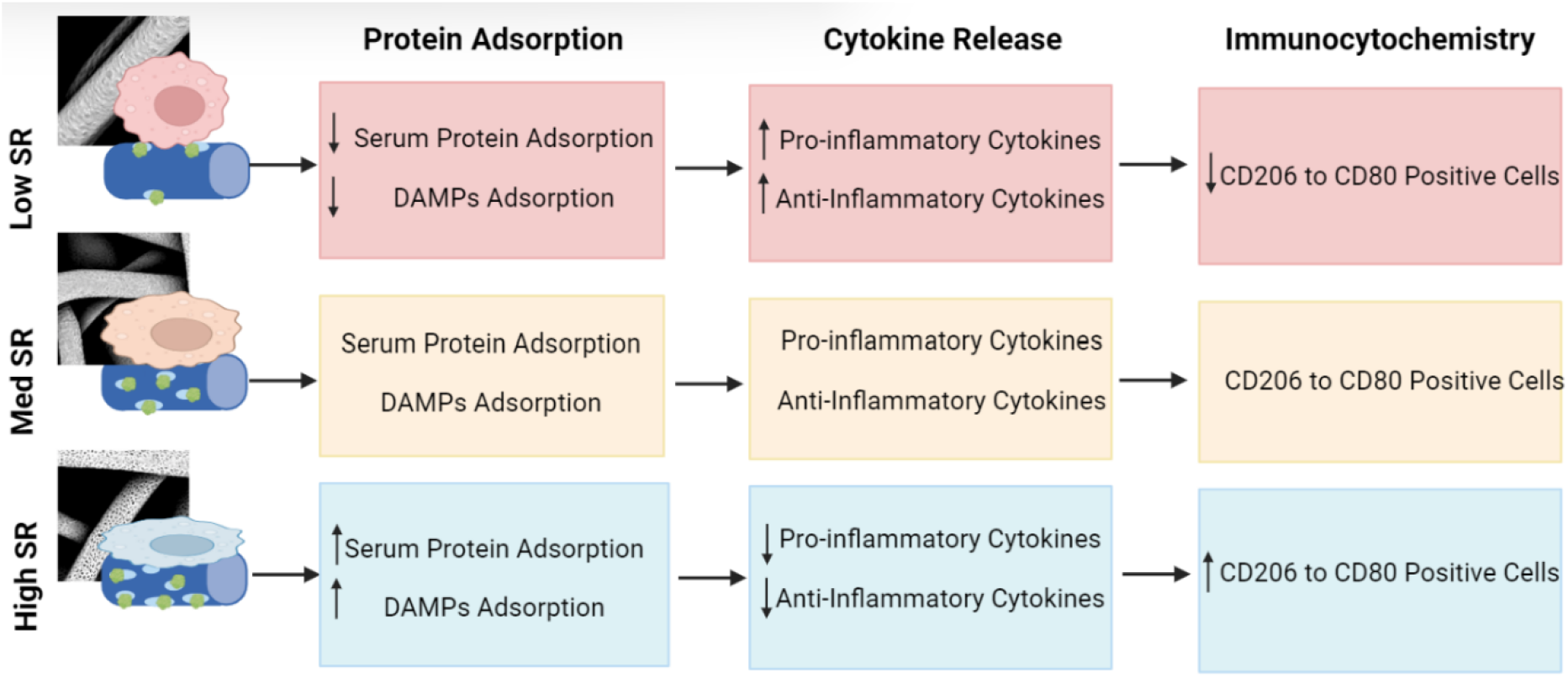

## 1. Introduction

Injuries to fibrous connective tissues suffer from poor intrinsic healing, leading to over 17 million doctor’s visits and 1.4 million surgeries annually in the US [1, 2]. Even when repair occurs, the resultant tissue has inferior mechanical properties, changing the biomechanics of the tissue and increasing the risk of reinjury [3-5]. After injury, inflammatory cytokines such as interleukin 1 (IL-1) and tumor necrosis factor alpha (TNF-α) are released, inhibiting healing and leading to the degeneration of tissues by promoting catabolic factors like matrix metalloproteinases (MMPs) and downregulating structural proteins like proteoglycans [6-12]. Current treatments often worsen this problem by promoting the release of damage-associated molecular patterns (DAMPs) and inflammatory signals [13, 14]. Inhibiting these inflammatory signals and promoting anti-inflammatory cytokines, like transforming growth factor beta (TGF-β), have been shown to improve healing in fibrous connective tissues [15-18].

The balance of pro-inflammatory and pro-regenerative factors is largely determined by the polarization of macrophages to either the pro-inflammatory (traditionally called M1) state or the pro-regenerative (M2) state [19-21]. While the M1/M2 dichotomy does not fully capture the dynamic nature of macrophage behavior, we have utilized it in this paper to remain consistent with prior literature [22, 23]. Persistence of macrophages with an M1-like phenotype has been shown to delay healing and produce less functional tissue [24-27], while a higher ratio of M2 to M1-like macrophages promotes healthy regeneration [28-30]. Depletion of macrophages in mice has been shown to accelerate degeneration of connective tissues, showing that this cell type is necessary for healing [31]. Thus, understanding the cues which drive macrophage polarization has been a major goal for researchers seeking to develop tissue regenerative therapies for fibrous connective tissue injuries [32, 33].

Biomaterials which mimic the structural properties of the native extracellular matrix (ECM) can promote early stages of healing, creating an opportunity to control macrophage phenotypes using engineered biomaterials [34-36]. Thus, research has focused on modulating biomaterial physical properties to control macrophage response in common 3D porous scaffolds used for tissue engineering, including hydrogels [37], collagen foams [38], 3D-printed scaffolds [39], and electrospun mesh [40]. Electrospinning is an advantageous fabrication technique based on its ability to fabricate nano-to micrometer sized fibers that mimic both the size and alignment of native collagen fibers. [41] Further, electrospinning has the ability to produce mesh with a large surface area to volume ratio, allowing for ample cell interaction [42].

Understanding how the properties of fibrous materials affect macrophage polarization and responses is a vital step in designing scaffolds which can manipulate the inflammatory microenvironment and promote regeneration. Properties such as fiber diameter [43-46], fiber alignment [47-51], porosity [52], and stiffness [53-60] have all been shown to affect macrophage phenotype. Studies in two-dimensions (2D) have found that surfaces with certain nanopatterns were shown to increase cell adhesion, cytokine release [61], cell migration [62], and phagocytic ability [63], while other patterns limited cell movement, decreased expression of pro-inflammatory cytokines [64, 65], and even promoted polarization to the M2 phenotype [66, 67]. Studies on 2D titanium surfaces found that smoother surfaces promoted a pro-inflammatory response, while rougher surfaces were anti-inflammatory [68] and produced greater amounts of TGF-β [69]. However, other studies showed on 2D titanium that rougher surfaces were pro-inflammatory [70] or that roughness had little effect [71]. A more recent study addressed this contradiction by showing that only a narrow range of roughness promoted an anti-inflammatory response in 2D, while surfaces that were either too smooth or too rough were pro-inflammatory [72]. Loss of the D-band, the major source of roughness on the collagen fiber with a period of around 67 nm [73], prevented attachment and migration of fibroblasts [74]. Surface roughness (SR) has been shown to modulate the behavior and differentiation of cells necessary for wound repair, including mesenchymal stromal cells (MSCs) [75-78]. The importance of controlling SR has been seen *in vivo*, as it has been demonstrated that increasing surface roughness of breast and dental implants leads to more inflammation and poorer outcomes [79, 80]. Finally, a recent study showed that increasing fiber SR decreased inflammation in mouse macrophages. However, this study used SR values much larger than that of the native ECM [81].

Adding nanostructures to materials can increase protein adsorption [82-84]. Materials with abundant binding sites for proteins like fibronectin can also activate the PI3K/AKT pathway and downregulate inflammatory pathways, including NF-κB [85]. Integrin binding to proteins adsorbed onto the material surface [86-90] promote cell growth and an elongated phenotype [85, 91, 92], associated with a shift in polarization towards an M2-like phenotype [93, 94]. Integrin binding to adsorbed proteins, including damage-associated molecular patterns (DAMPs), triggers formation of the focal adhesion complex [95], leading to recruitment of adhesion adaptor proteins, such as paxillin and focal adhesion kinase (FAK), that strengthen the focal adhesion [96]. These have been shown to be essential for cell adhesion, migration [97, 98] and inflammatory signaling [99]. Conversely, blocking integrin signaling prevents ECM binding and inflammatory activation [100-103].

In this study, the impact of fiber SR on macrophage polarization was assessed both *in vitro* and *in vivo*. Mesh with increasing SR were fabricated by modulating vapor-induced phase separation during the electrospinning process. Adsorption of serum proteins and DAMPs was assessed using a BCA assay. Polarization of human monocyte-derived macrophages from three donors was assessed via cytokine release at multiple timepoints and immunostaining for phenotype-specific cell surface markers after five days. *In vivo*, mesh of various SR were implanted subcutaneously in mice for three days to assess macrophage polarization using the same cell surface markers utilized in the *in vitro* studies. These studies provided important insight into the physical cues which drive macrophage polarization and how these signals can be intentionally manipulated to promote a regenerative phenotype.

## 2. Materials and Methods

### 2.1 Scaffold Fabrication and Characterization

Microfibrous mesh were fabricated using an in-house electrospinning setup. A 20% (w/v) solution of 50,000 kDa polycaprolactone (PCL) in chloroform was pumped through a 21-gauge needle tip at a rate of 1.5 mL/hr into an electric field generated using a voltage source and a grounded copper plate distanced 30 cm from the needle tip. Ambient relative humidity (RH, 30%, 50% and 70%) was controlled during the electrospinning process to promote vapor-induced phase separation (VIPS). The accelerating voltage was adjusted to maintain equal fiber diameters between groups with 12kV used for the 30% and 50% RH and 15kV used for the 70%. The mesh were then left under a fume hood overnight to allow any remaining chloroform to evaporate.

To promote electron conductivity, 8 mm punches of mesh were taken and sputter-coated with 5 nm of gold using a Q150T sputter coater (Quorum Technologies). Morphological analysis for fiber alignment, fiber diameter, percent porosity, and fiber surface texture was carried out using a Phenom scanning electron microscope (SEM) with five images taken from three mesh for each humidity group (n=15). Fiber diameter was analyzed using the DiameterJ plugin in ImageJ.

To measure surface roughness, 8 mm punches were analyzed using an ICON atomic force microscope (AFM; Bruker) in tapping mode. To determine the surface roughness, five 1 μm x 1 μm sections of fiber surface were analyzed from three mesh of each humidity group (n=15).

### 2.2 Quantification of Protein Adsorption on Mesh by BCA

The impact of SR on protein adsorption was assessed using both fetal bovine serum (FBS) and DAMPs. Murine 3T3 fibroblasts (5 million), suspended in 10 mL of phosphate-buffered saline (PBS) were lysed via repeated freeze-thaw cycles to create a source of DAMPs. Eight mm punches of mesh were incubated in 300 μL of either DAMP solution or 40% FBS at 37 °C for 24 hours. Afterwards, the mesh were washed three times in PBS to rinse off any protein that had not yet adsorbed, then incubated in 100 μL of 0.1% SDS twice to strip the adsorbed protein off the mesh. The SDS-protein mixture was measured for total protein content using a bicinchoninic acid (BCA) assay (ThermoFisher) which was analyzed for absorbance at a 562nm wavelength using an Infinite M200 Pro plater reader (Tecan). In each experiment, five punches were used from three different mesh (n=15).

### 2.3 Monocyte Isolation and Cell Seeding

Whole blood samples were drawn from three healthy male volunteers, aged 20-35 (University of Washington IRB #00018882 accepted November 28, 2023). All donors were provided and signed informed consent documents. Monocytes were separated from whole blood using a Rosette-Sep Human Monocyte Enrichment Cocktail (Stem Cell Technologies) combined with Lymphoprep (Stem Cell Technologies) density gradient in a Sep-Tube (Stem Cell Technologies). Monocytes were cultured for 5 days in RPMI media supplemented with 10 ng/mL of macrophage colony stimulating factor (MCSF) to differentiate them into macrophages.

The night prior to cell seeding, 12 mm punches of mesh were serially incubated in 70% ethanol, 50% ethanol, 30% ethanol, and DI water for 30 minutes each to promote aqueous solution penetration, fully hydrate the mesh, and sterilize. The mesh were then incubated overnight in RPMI media supplemented with 40% FBS to ensure ample protein adsorption. The next day, macrophages were seeded onto the prepared mesh at a density of 60,000 cells/mesh.

### 2.4 Cytokine/Chemokine Quantification via Multiplex Immunofluorescent Assay

Macrophage phenotype was assessed by measuring the release of pro- and anti-inflammatory cytokines over a 120-hour period. Monocytes were incubated at 37 °C, 5% CO_2_ for five days in a 24-well plate with three punches from three mesh (n=9 per donor, n=27 total) used per humidity group (RH, 30%, 50% and 70%). Cultured media was sampled (10% total volume in the well) at 3 hours, 24 hours, 72 hours, and 120 hours. Released pro- and anti-inflammatory cytokines were then measured in these samples using a Luminex assay (Procartaplex) for pro-(interleukin 1-beta (IL-1β), interleukin 23 (IL-23), tumor necrosis factor alpha (TNF-α), and regulated on activation normal T-cell expressed and secreted (RANTES)) and anti-inflammatory (interleukin 10 (IL-10), latency associated protein (LAP), interleukin 13 (IL-13), and interleukin 4 (IL-4)) cytokines run on a Bio-Plex 200 reader (BioRad). Data was collected from only two donors for RANTES, LAP, and IL-13.

### 2.5 Immunofluorescence Staining for Cell Surface Markers

Macrophage phenotype was assessed by immunofluorescence (IF) staining for cell surface markers specific to the M1 and M2 phenotypes. CD80 was used as an M1 marker [104] and CD206 was used as an M2 marker [105]. Monocytes from each donor were incubated on the mesh at 37 °C for five days in a 24-well non-treated plate with three punches from three mesh (n=9 per donor, n=27 total) used per humidity group. After five days, the mesh were fixed in 4% paraformaldehyde for 15 min. The mesh were then blocked in 2% BSA for one hour at RT prior to immunostaining for CD80 (Invitrogen #MA5-15512; diluted 1:100) and CD 206 (Invitrogen # MA5-32498; diluted 1:200). Samples were incubated in primary antibodies overnight at 4 °C and in secondary antibodies (Invitrogen #A11036 and #A32728, diluted 1:1000) for 2 hours at RT. Fluorescent images were then taken using a Leica SP8 confocal microscope. Images were false colored for consistency and visual clarity.

### 2.6 Subcutaneous Implantation in Mice and Immunohistochemistry

The impact of mesh SR on macrophage polarization was assessed *in vivo* using a mouse model with fluorescently labeled macrophages. Mesh from each humidity group were implanted on the dorsum of 4 month-old MacGreen (B6.Cg-Tg(Csf1r-EGFP)1Hume/J; the Jackson Laboratory #018549; 3 male, 1 female) mice (University of Kansas Medical Center IUCAC 23-01-294 approved September 9, 2023). Mesh from each humidity group, as well as a PCL film control, were implanted in each mouse (4 mesh total per mouse), with four mice used (n=4). Location of each mesh was randomized to minimize location-specific effects. After three days, the mice were sacrificed, mesh were recovered, were fixed overnight in 4% paraformaldehyde (PFA), washed in PBS, and placed in a 30% sucrose solution for 48 hours. Samples were then flash frozen in OCT Compound (Fisher) using liquid nitrogen and sectioned. Sections were then immunostained and imaged using the protocol above. Images were false colored for consistency and visual clarity.

### 2.7 Statistical Analysis

Results were tested for normality using an Anderson-Darling test and for equal variance using a Brown-Forsythe test. If both these conditions are met, then results were compared statistically using a one (condition or time effects) or two-way (condition and time effects) ANOVA with a Bonferroni post hoc test. If not, the non-parametric Kruskal-Wallis test was used. Differences were considered statistically significant for p-values below 0.05. All analyses were performed using Graph-Pad Prism (GraphPad Software). Sample sizes were determined using a power analysis (G*Power) with α = 0.05 and 1-β = 0.80 and all studies met the criteria determined by these tests.

## 3. Results

### 3.1 Scaffold Characterization

The impact of increasing RH on fiber morphology and SR was assessed visually via SEM and quantified via AFM. Analysis of SEM micrographs demonstrated a visual increase in roughness on the surface of electrospun fibers with increasing RH (**Figure 1A**). Furthermore, there was no statistically significant difference in fiber diameter or percent porosity between humidity groups (**Figure 1B)**. AFM analysis showed a visual increase in surface topography with increasing RH (**Figure 1C**). Further, quantification of the arithmetic average roughness (Ra) and surface area showed a statistically significant increase in roughness and surface area with increasing RH. Humidities of 30%, 50%, and 70% corresponded to Ra values of 11.0 ± 4.2, 28.9 ± 7.8, and 60.6 ± 10.4 nm respectively (**Figure 1D)**.

**Figure 1:**
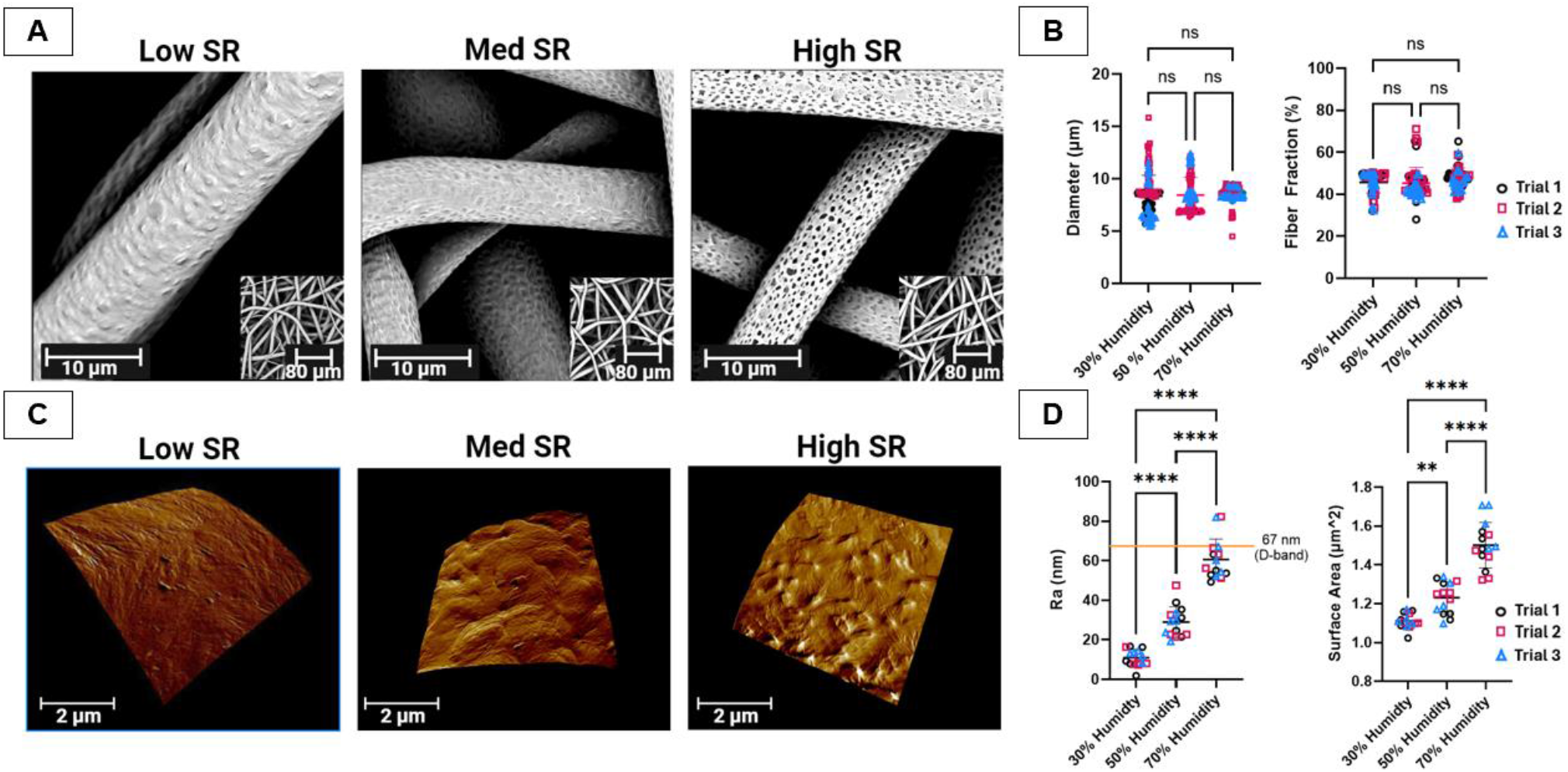
Modification of Fiber SR with Increasing RH Without Significant Change to Diameter or Porosity. Electrospun PCL were imaged with SEM (A). Fiber diameter and porosity (B) were then determined and compared. Fiber surface roughness (SR) was visualized and quantified using AFM (C) *p<0.05, ***p<0.001, ****p<0.0001

### 3.2 Protein Adsorption

The impact of increasing SR on the absorption of proteins was analyzed using a BCA assay. There was a statistically significant increase in the adsorption of FBS proteins with increasing SR. Adsorbed protein amounts on the low, medium, and high SR mesh were 22.9 ± 11.2 μg, 48.8 ± 26.7 μg, 72.5 ± 33.5 μg respectively (**Figure 2A**). There was a statistically significant increase in DAMP adsorption on the high SR mesh, but differences between the low and medium SR mesh were non-significant. Adsorbed DAMP protein on the low, medium, and high SR mesh were 17.3 ± 10.2 μg, 11.2 ± 9.0 μg, 28.3 ± 16.7 μg respectively (**Figure 2B**).

**Figure 2:**
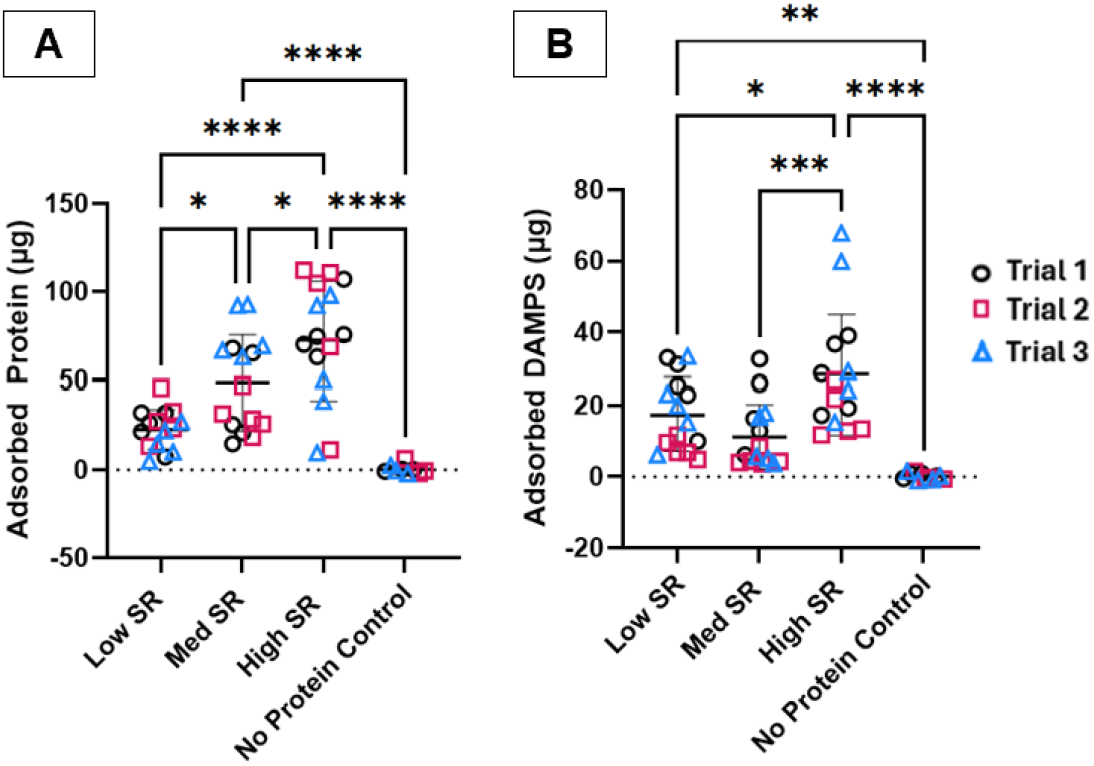
Protein Adsorption on Mesh. Mesh were incubated in 40% FBS (A) or a DAMP solution (B). Adsorbed protein was quantified via a BCA assay. * p ≤ 0.05; **p≤ 0.01: *** p ≤ 0.001; **** p ≤ 0.0001

### 3.3 Cytokine Release

The release of pro- and anti-inflammatory cytokines from primary human macrophages cultured on mesh with increasing SR was analyzed using an eight-plex Luminex assay. Results for pro- and anti-inflammatory cytokines are presented in aggregate **(Figure 3A and 4A respectively)** and at the 24- and 72-hour timepoints **(Figure 3B and 4B respectively)**. Results were analyzed by two- and one-way ANOVA respectively. In general, cytokine release peaked at 24 hours and showed a reduction in both pro- and anti-inflammatory signaling with increasing SR. The pro-inflammatory cytokines IL-1β and TNF-α and pro-inflammatory chemokine, RANTES, showed a statistically significant increase comparing low to high SR at 24 hours. Levels of IL-1β and RANTES remained elevated in the low SR group compared to the high SR group at 72 hours. Expression of IL-23 was elevated in the medium SR group at 72 hours, but this was largely driven by results from a single donor. The expression of the anti-inflammatory cytokines IL-10 and LAP, a cleavage product of activated TGF-β, also showed a statistically significant increase in the low SR group at 24 hours. Expression of LAP and IL-13 were elevated in the medium SR group at 72 hours, driven by results from a single donor. Overall, expression of both pro- and anti-inflammatory cytokines was increased on the low SR mesh.

**Figure 3:**
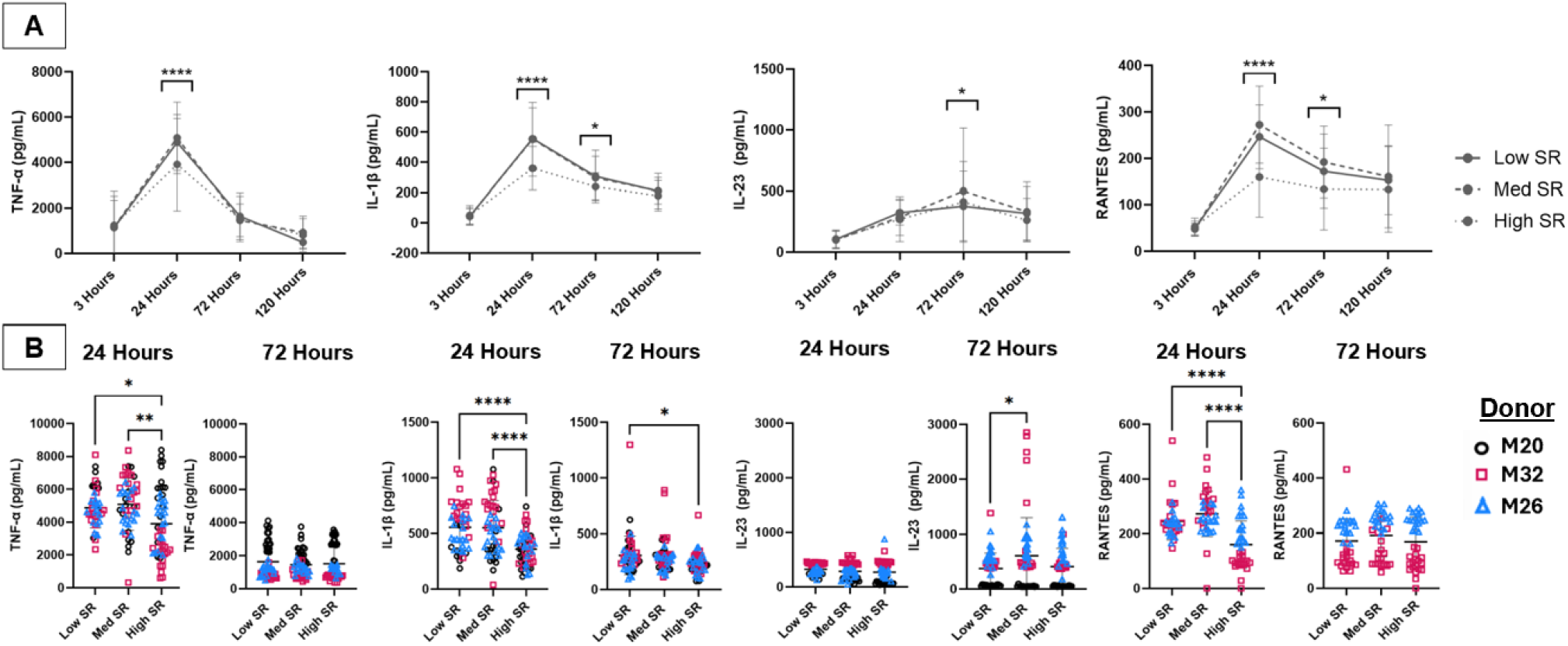
Pro-Inflammatory Cytokine Release. Cultured media from primary macrophages was tested on an eight-plex Luminex assay at 3, 24, 72, and 120 hours for pro-inflammatory cytokines. Individual figures for 24- and 72-hour timepoints * p ≤ 0.05; **p≤ 0.01: *** p ≤ 0.001; **** p ≤ 0.0001

**Figure 4:**
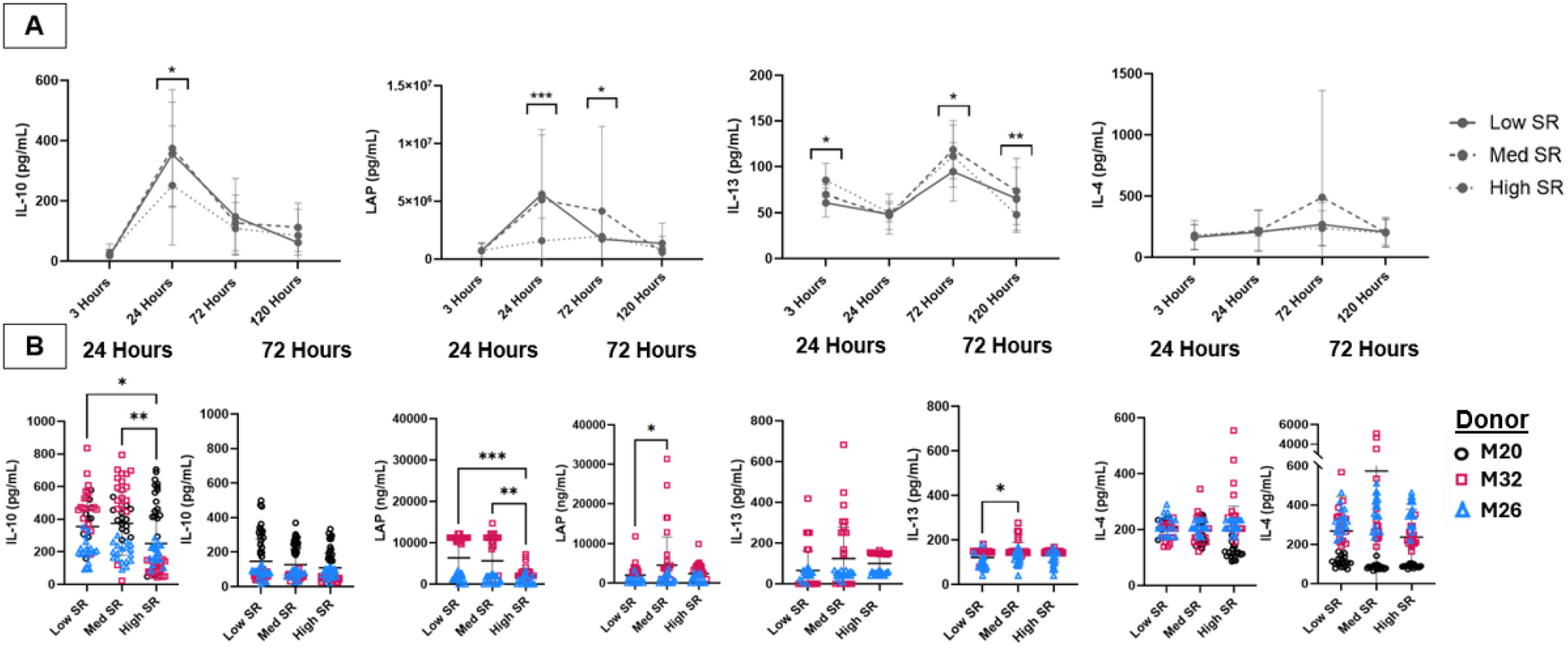
Anti-Inflammatory Cytokine Release. Cultured media from primary macrophages was tested on an eight-plex Luminex assay at 3, 24, 72, and 120 hours for anti-inflammatory cytokines. Individual figures for 24- and 72-hour timepoints * p ≤ 0.05; **p≤ 0.01: *** p ≤ 0.001; **** p ≤ 0.0001

### 3.4 Immunostaining

The expression of cell surface markers associated with the M1 and M2 macrophage phenotypes on *in vitro* samples was assessed using ICC staining for CD80 (M1) and CD206 (M2). This showed a visual increase in staining for CD206 with an increase in SR **(Figure 5A)**. ImageJ was used to count the number of cells positively stained for CD80 and CD206. Counts were then normalized to total cell number and compared to each other to determine the M2 to M1 ratio. This showed a statistically significant increase in the ratio of staining for CD206 to CD80 with increasing SR **(Figure 5B)**.

**Figure 5:**
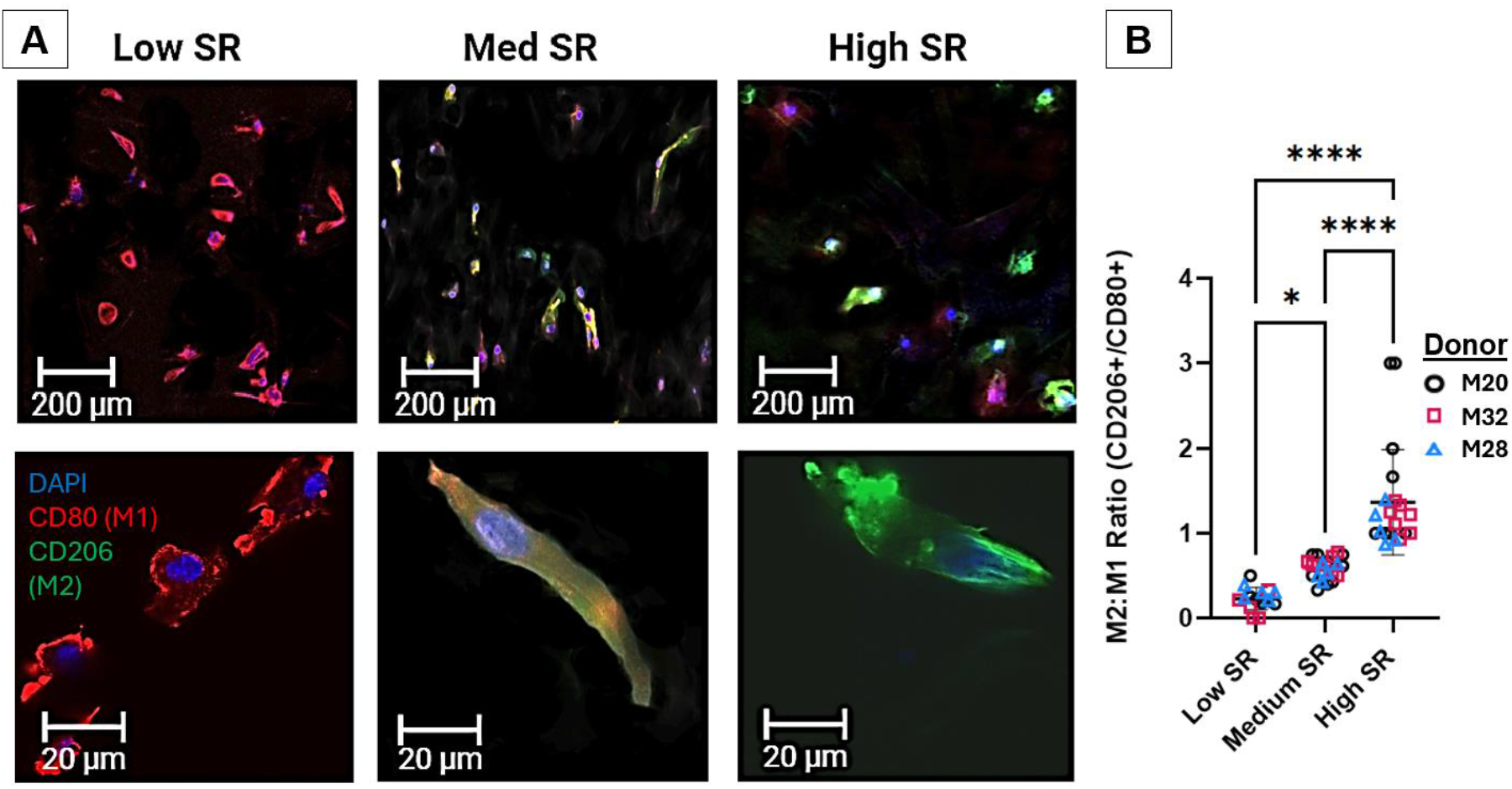
Immunocytochemistry Staining of Primary Macrophages for CD 80 (M1) and CD 206 (M2). Images of CD 80 and 206 expression in primary macrophages incubated on mesh with increasing SR. * p ≤ 0.05; **** p ≤ 0.0001. Images were false colored for consistency and visual clarity.

### 3.5 *In vivo* Mouse data

The polarization of macrophages *in* vivo after subcutaneous implantation of electrospun mesh with increasing SR in MacGreen mice was assessed using Immunohistochemistry (IHC) staining for CD80 (M1) and CD206 (M2). A schematic view of fiber sections and macrophage infiltration is seen in **Figure 6A**. Overall macrophage morphology and density on fibers is seen in **Figure 6B**. GFP-labeling of macrophages showed that immune cell infiltration was consistent between groups **(Figure 6C and E)**. A visual increase in staining for CD206 was observed with increasing SR. However, staining for CD80 remained consistent between samples **(Figure 6E)**. Quantification, using the protocol above in ImageJ, showed a statistically significant increase in the ratio of cell staining for CD206 to CD80 with increasing SR. All cells on the flat film control stained positive for CD80 and negative for CD206 **(Figure 6F)**.

**Figure 6:**
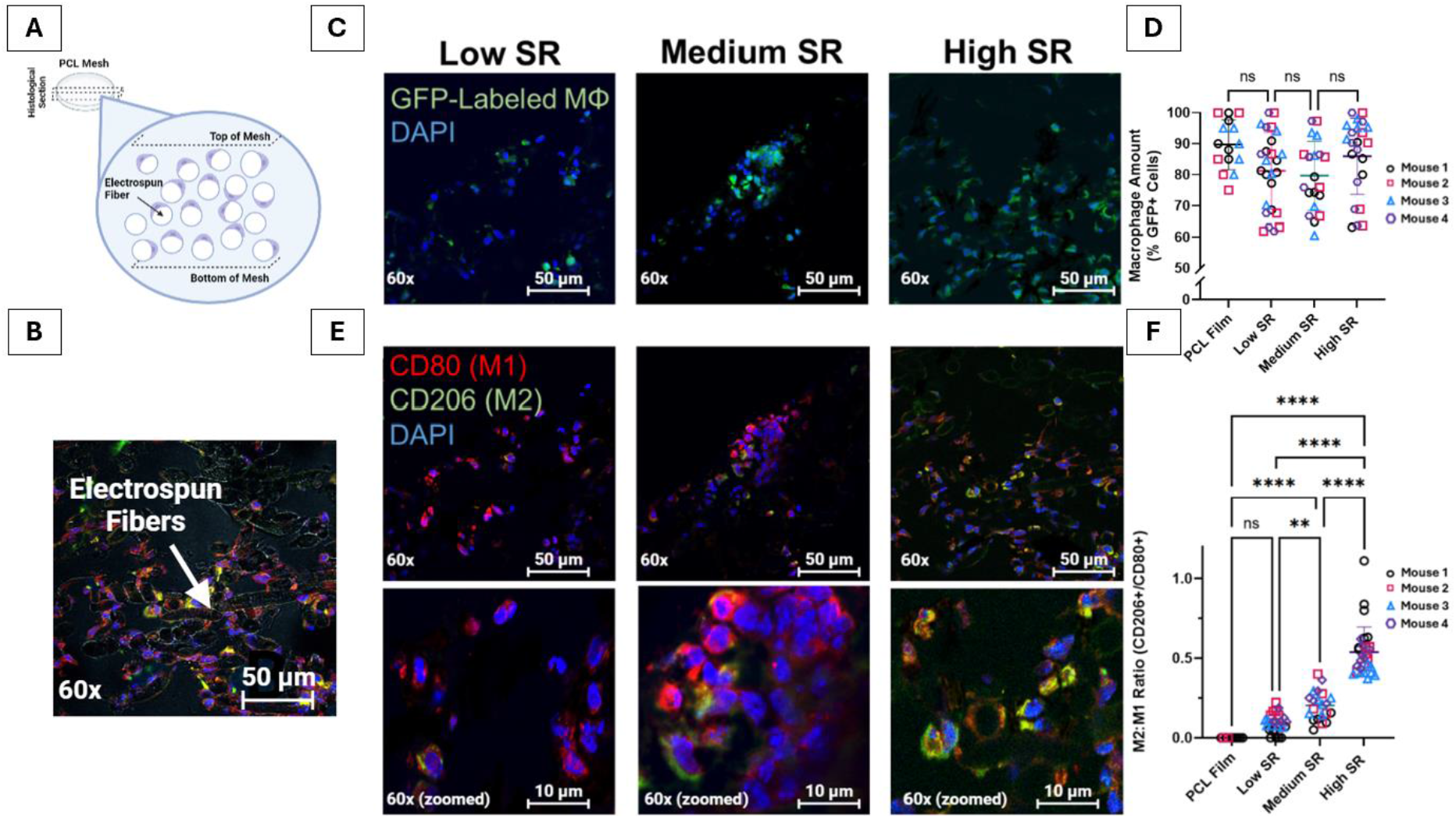
Immunohistochemistry Staining of Mesh after Implantation in MacGreen Mice for CD80 (M1) and CD206 (M2). Schematic (A) and actual (B) visualization of macrophage morphology on fibers. Infiltration (C and D) and polarization (E and F) of macrophages imaged at 60x after subcutaneous implantation in mice. Cropped and zoomed in images of B are shown in the lowest panels. A brightfield image shows interaction between cells and mesh (C). Ratio of cells expressing CD206 and CD80 was measured in ImageJ (D). ** p ≤ 0.01; ****p<0.0001. Images were false colored for consistency and visual clarity.

## 4. Discussion

These findings show that macrophage phenotype can be modulated by modifying the surface of micron-sized fibers used for tissue engineering. This provides valuable information that will help guide the design of future biomaterials intended to promote regeneration.

By modulating RH from spinning, fiber with Ra values of 11.0 ± 4.2 nm to 60.6 ± 10.4 nm were produced. This corresponds with results from Chen, et al., who used RH values from 20% to 70% to produce fibers with Ra values from 4.3 ± 2.5 nm to 71.0 ± 11.0 nm [78]. Adsorption of serum proteins increased 3.2 times with the increase in SR from the low to high group, corresponding to other results showing a 3 to 4 times increase in protein adsorption with increasing SR [78, 82]. Further studies are necessary to identify the chemistry and conformation of specific proteins which adsorbed and to interrogate how they affect important pathways such as PI3K/AKT.

Other researchers have utilized nanoscale roughness on fibrous biomaterials to promote regenerative phenotypes in other cell types, such as mesenchymal stromal cells [78]. Zhu, et al. explored the impact of fiber roughness on macrophages, but they used a different technique to create SR in their system, used much larger SR values, and used mouse RAW 267.4 macrophages for their *in vitro* studies [81]. They found that adding shish-kebab structure to their fibers decreased release of pro-inflammatory cytokines and increased the M2:M1 ratio *in vivo* in mice, corresponding with our results. However, they saw an increase in anti-inflammatory cytokine release in their rough samples, which was not seen in our system. This was the first study to show the impact of fiber surface roughness on the scale of native ECM using primary human macrophages and adds to the body of research on how electrospun fiber properties impact macrophage polarization and can be leveraged to promote regeneration.

Our results showed that pro-inflammatory signaling peaked at 24 hours and was attenuated after 72 hours. This corresponds to the natural inflammatory response after injury in which monocytes infiltrate into the wound environment, differentiate to the M1 state, clear pathogens and dead tissue, then differentiate again to a pro-regenerative M2 state after approximately three days [106-108]. This corresponds with results from Saino, et al. illustrating that cytokine release on randomly aligned, micron-sized fibers was higher at 1 day than at 7 days [43]. However, studies on other systems, such as biotinylated bone scaffolds, saw cytokine release peak at later time points [109].

Our studies found that increasing fiber SR led to a decrease in the release of pro-inflammatory cytokines including TNF-α and IL-1β. Inhibition of these signals has been shown to improve healing in fibrous connective tissues [17]. Further, our ICC data showed an increase in the M2:M1 ratio with increasing SR, a change that has also been shown to improve healing [28]. Studies on other *in vitro* biomaterial systems have shown that shifts towards a more M2-like macrophage phenotype can regulate ECM assembly [110]. Abaricia et al. and Barr, et al. both found that increasing SR on implants led to increased inflammation and worse outcomes *in vivo* [79, 80]. Our results support the idea that there is a narrow band of SR values which decreases the inflammatory response. This information will be useful for future implant design.

Cells in our system showed a decrease in both pro- and anti-inflammatory cytokine and chemokine release with increasing SR. However, our ICC data did show an increase in M2-associated cell surface markers with increasing SR. This difference may be explained by a temporal lag between cells changing phenotype and releasing anti-inflammatory cytokines. The contribution of other immune cells in resolving inflammation, not currently modeled in our system, may provide additional information. It is also possible that staining for additional M1 and M2-associated cell surface markers would help to explain these inconsistencies. Finally, Spiller, et al. found that on porous collagen sponges, the release of anti-inflammatory cytokines was actually highest in M1 polarized macrophages, showing that the traditional paradigms of macrophage phenotype and cytokine release cannot always be relied upon [111].

These studies were limited by their reliance on the M1/M2 dichotomy, which cannot adequately describe the full spectrum of macrophage behavior. Further studies are necessary to fully characterize the cellular response to increasing SR. Further, our immunofluorescence staining only assessed the cells at a single time point. Further studies are needed to assess the dynamic response of macrophages to increasing SR. Further studies are also required to determine the exact range of SR values which produce an M2-like response in macrophages, as the top SR achievable in our system was limited. Finally, studies are needed to determine if the same effect on macrophage polarization is seen on fibers with nano-scale diameters.

## 5. Conclusion

In summary, an electrospun fiber scaffold system with tunable surface roughness mimicking ECM structure was produced. Increasing fiber SR led to increased protein adsorption and, subsequently, changes in the polarization of macrophages which were incubated on these materials. Increasing the SR led to a decrease the expression of pro-inflammatory cytokines which are known to play a role in the degeneration of fibrous connective tissues and to increase the M2:M1 ratio, which is known to improve healing. Overall, tuning the SR of electrospun fibers shows promise as a tool to increase the regenerative potential of fibrous biomaterials used to improve healing after fibrous connective tissue injuries.

## Acknowledgements

Funding from University of Washington Startup and Endowment, DK126456, DK127497, and DK131716. Support from the UW Molecular Analysis Facility, and the Lynn and Mike Garvey Imaging Core. Images were created in Biorender.com.

